# Rapid, Affordable, Collection and Analysis of Bioaerosol Viral Pathogen (Begomoviruses and Whitefly Vectors)

**DOI:** 10.1101/451559

**Authors:** Wayne B. Hunter, Cindy L. McKenzie, Bailey W. Mitchell

## Abstract

Detection of pathogens is critical to monitoring their distribution and spread, and is a key component in the prediction and management of disease epidemiology. Monitoring for pathogens as bioaerosols requires developing techniques which are sensitive, affordable, and time saving before they will have widespread impact. This approach also overcomes private property issues, which are a major pitfall in monitoring diseases in complex agricultural and urban settings. In this study, we have applied an emerging technology of electrostatic sampling to the detection of an insect-transmitted plant pathogen as a bioaerosol. Where insects aggregate in large numbers, as with whiteflies, leafhoppers, psyllids and honey bees, the pathogen (ie. virus or bacteria) becoming aerosolized as thousands of excreta droplets fall from the plants during feeding. Agricultural systems have not fully measured the impact of bioaerosols on disease epidemiology. Electrostatic sampling provides a valuable, affordable, method for monitoring for diseases as bioaerosols, which includes plant, animal and human pathogens. This study shows results which successfully used an electrostatic sampling device to collect an aerosolized begomovirus from the air near whiteflies feeding on virus-infected tomato plants.

## INTRODUCTION

Today’s political climate of uncertainty contains warnings of dangers from biological agents that may cause harm to humans, animals, or food crops. These concerns include agents that can be spread as airborne particulates and have spurred renewed interest in the development and application of methods to collect, sample, and identify airborne organisms and chemicals. Microorganisms such as bacteria, fungal spores, and viruses carried by air movements are referred to as bioaerosols (Harper 1961; Killingley et al, 2013; Kumar et al, 2018). While most research on bioaerosol sampling has focused primarily on human and animal pathogens (Belser et al, 2013; 2016; Gustin et al, 2013; Kormunth et al, 2018; Mitchell et al, 2003; Roux et al, 2013; Suresh and Ricke 2002; Yamamoto et al, 2010). This study applied this emerging technology to the detection of insect-transmitted plant pathogens, specifically the whitefly-transmitted Begomoviruses that threaten many of the subsistence crops worldwide (Czosnek et al, 2017; Moffat 1999: Polston and Andersen 1997: Rosario, et al, 2014). Some of the advantages being realized are the ability to capture and detect diseases as bioaerosols, thus reducing the time and costs associated with monitoring disease presences or spread (Roux et al, 2013). Reducing costs and time in detection are critical components in the management of viral pathogens in humans (Cowling et al, 2013; Kormunth et al, 2018; Paynter 2015), but also when trying to manage and detect pathogens in urban or agricultural settings (Brodie et al, 2007). Electrostatic sampling is a rapid method to screen for the occurrence of human, animal, or plant pathogens which is not restricted by private property lines, crop type, or variety of samples collected, as the air flow provides a broad ‘sampling stream’ (Holt et al, 1999; Parvaneh, et al, 2000; Mehta et al, 2000; Suresh and Ricke 2002).

Recent advances in sampling, combined with breakthroughs in molecular techniques, and bioinformatics, permits the rapid collection and identification of microbes, like virus and bacteria, in a viable condition (Agranovski et al, 2005; Gerone et al, 1996). Thus, the microbe may be cultured, as well as extracted using the DNA and/or RNA sequences to identify each organism. (Hunter et al, 2001; Funk et al, 2001; Hunter and Polston 2001; Marutani-Hert et al, 2009; Rosario, et al, 2014; Sinisterra et al, 2005; Valles et al, 2004, 2008; 2018). However, genetic analyses works independently of the ability to propagate a microbe in culture (Chen et al, 2016; Hunter et al, 2003, 2009; Tokarz et al, 2018; Valles et al, 2018). The combination of these techniques thus have propelled the application of bioaerosol sampling into use as a new and rapidly evolving field of pathogen detection. The use of Electrostatic Sampling Devices (ESD) provides advantages that are beyond the normal detection methods of sample collecting. The microbe can be collected on a grounded piece of metal, or directly into selective or general culture media (when sampling for bacteria or fungi) (Gast et al, 1999; Holt et al, 1999; Mitchell et al, 2002; Willeke and Macher, 1999; Richardson et al, 2003ab). Recognizing the future of bioaerosol sampling this study evaluated if the small, newly designed electrostatic sampling device, ESD, would be a suitable to collect enough virus as bioaerosol for detection and monitoring of insect-transmitted plant pathogens. Air samplers are designed to monitor buildings which use continuous air sampling, the needs of agricultural cropping systems are focused on a device that is low in cost, easy to transport and sterilize, while being adaptable for a variety of agricultural crops.

The Electrostatic Sampling Device (ESD) designed by Drs. B. Mitchell and R. Gast (Patent 7,046,011, May 2006) was specifically designed to collect airborne particulates such as bacteria, viruses and fungal spores by using sharp-pointed electrodes that produce a focused electrostatic field with a high negative, direct-current voltage, generating a strong electrostatic field close to a grounded collector (FIG. 2A). The grounded collector is typically an agar plate with selective media on it for bacterial collections, or a dry metal plate, disc or rod to collect viruses. As charged particles are drawn to the collector, air adjacent to the particles is also drawn in, causing an airflow stream directed toward the collector (FIG. 2B). The number of microorganisms collected by this method is equivalent to at least that of a 100 L/min forced air impaction sampler (Gast et al, 2004; Mitchell et al, 2003). The numbers of organisms collected by the ESD in a given unit of time was very consistent in repeated trials collecting airborne bacteria, *Salmonella* and *Enterobacteriacea*, from poultry houses, produced deposition patterns of similar uniformity to those obtained by settling plates or impaction samplers (Gast et al, 2003, 2004; Mitchell et al, 2003; Richardson et al, 2003b). The ESD has an equivalent airflow rate of approximately 100 Liters/min, and the collected organisms are tightly held by electrostatic attraction. The airflow patterns have been demonstrated artificially using smoke sticks adjacent to the sampler, and in real life direct comparisons between impaction samplers and the ESD have shown equivalent results (Gast et al, 2003, 2004; Richardson et al, 2003b) (Fig. 2B). The ESD has no blower, the sampling area is not limited to a chamber, it requires a very low power source, thus enabling it to operate for 16 hrs on two small 9V batteries, The ESD is inexpensive to manufacture, rugged, and easy to disinfect (Mitchell et al, 2003).

**Fig. 1.**
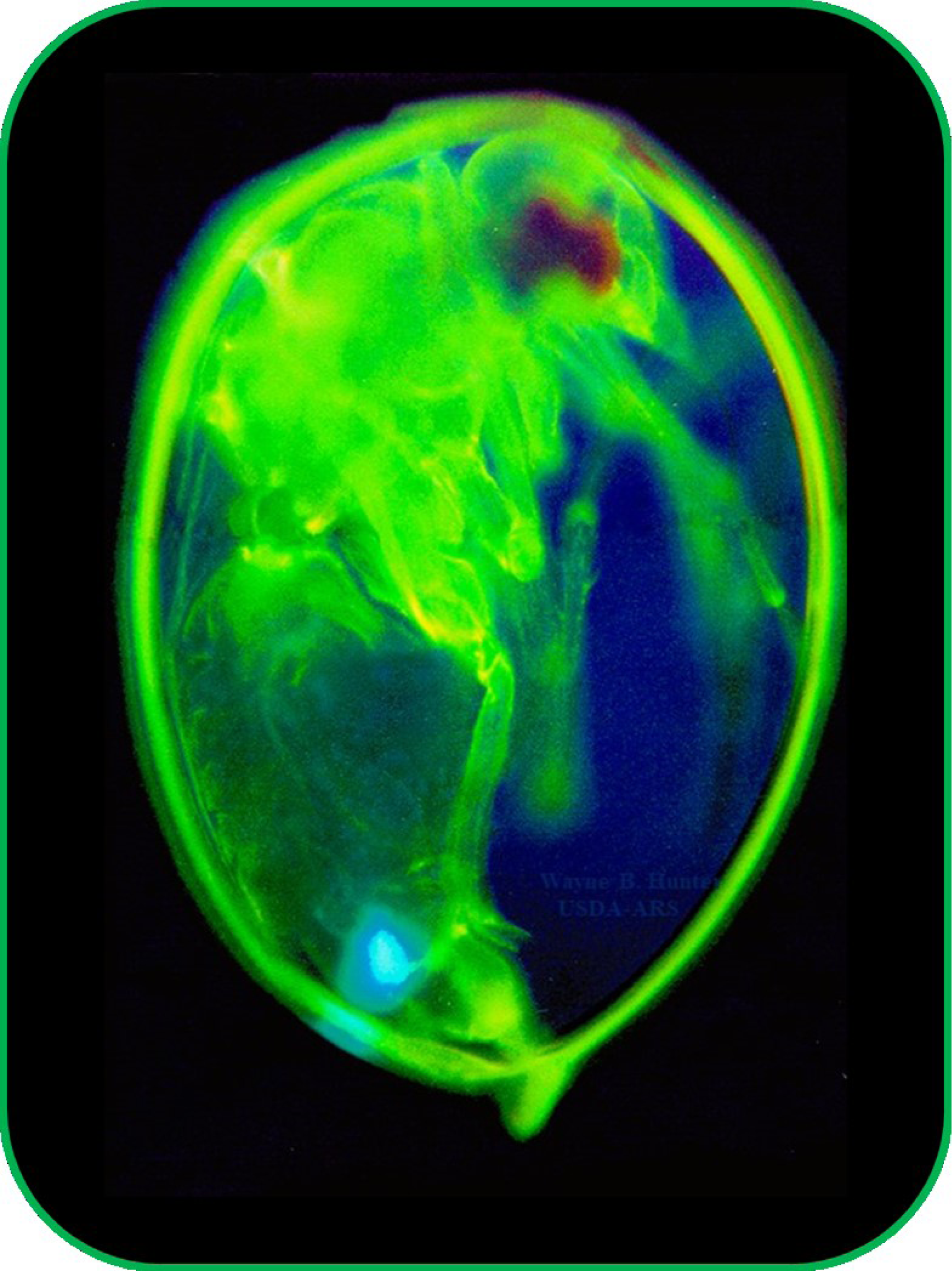
Immunostained Whitefly, *Bemisia argentifolii* (aka. *B. tabaci,* B-Biotype) (Hemiptera: Aleyroididae) Vector of Begomoviruses (Photo: Wayne Hunter, USDA-ARS).

**Figure 2.**
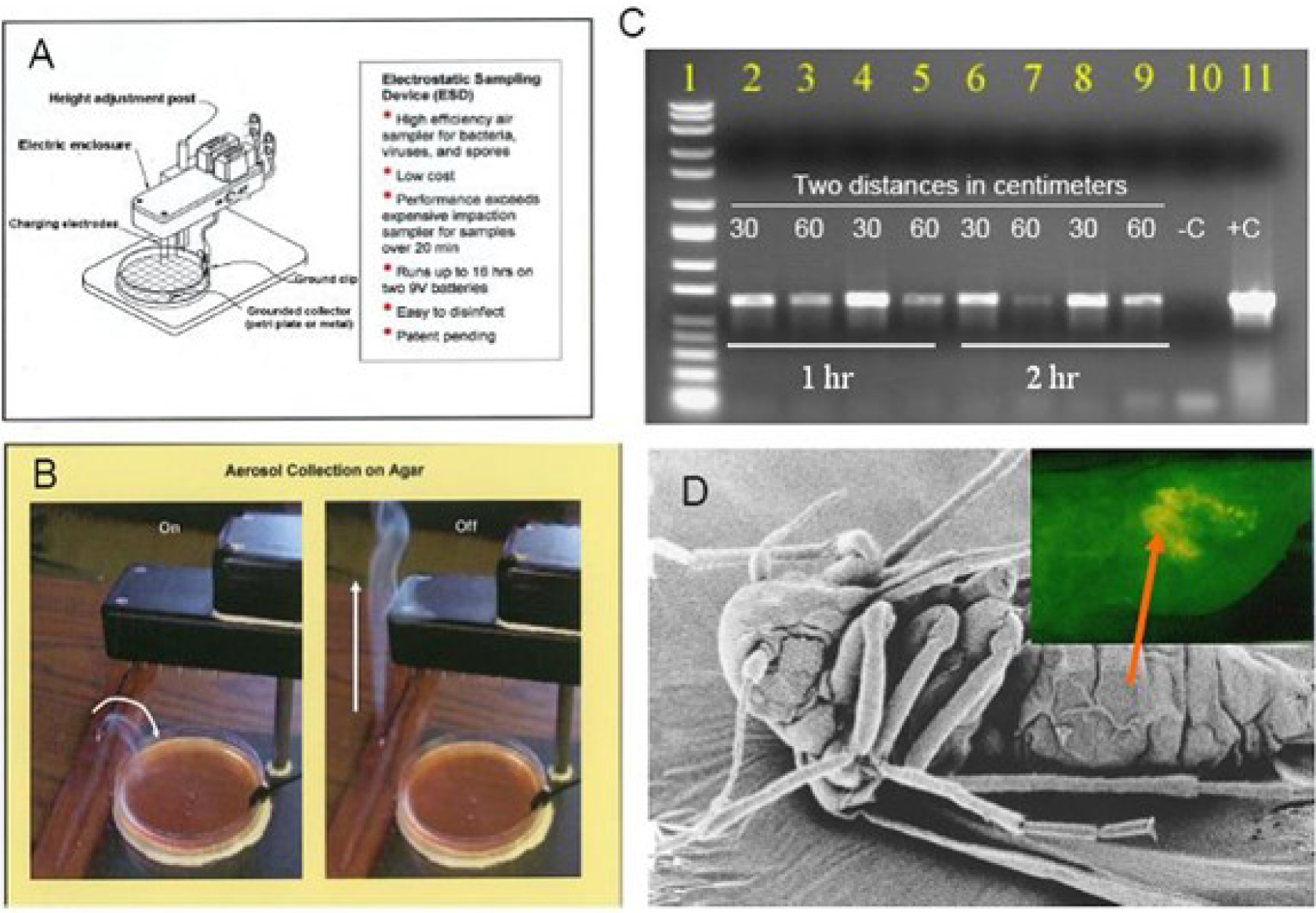
Capture, collection and detection of the Begomovirus, Tomato yellow leaf-curl virus, TYLCV, in air as a bioaerosol using an electrostatic sampling device, ESD. (**A**) Diagram of electrostatic device, ESD. (**B**) Smoke used to show electrostatic pull of bio-aerosols a grounded agar plate. The collecting surface may also be a metallic plate, rod, or agar plate (Mitchell et al, 2002; Mitchell and Gast 2003). (**C**) Gel showing results of samples collected from the grounded metal scoopula, sterile cotton swab, when ESD was placed at two distances, 30 and 60 cm, from tomato plant infested with *B. tabaci*. Durations were one and two hours respectively. Begomovirus source was an uncaged whitefly colony feeding on a TYLCV infected tomato plant. (**D**) Scanning electron micrograph image of a whitefly adult, with inset showing FITC immunolabeled TYLCV within the whitefly digestive system after feeding on infected tomato (arrow)(method in Hunter et al, 1998). TYLCV enters the hemolymph and replicates within the whitefly (Sinisterra et al, 2005). Virus passes out with saliva (Rosell et al, 1999) and excreta during feeding becoming aerosolized.

Begomoviruses are whitefly-transmitted, single-stranded, DNA plant infecting viruses (Czosnek et al, 2001; Polston and Andersen 1997; Zerbini et al, 2017), and are known for reduce major food crops world-wide, including tomato, beans, and cassava (Njorage, et al, 2017; Moffat 1999; Saeed and Samad 2017). Virions leave the insect body either *via* the salivary ducts, passing down the insect’s stylets during feeding thus being deposited into plant tissues (Ghanim et al, 2001; Rosell et al, 1999) or along with excreta as droplets into the air. Excretion of virus is most likely the route of virus becoming aerosolized (Czosnek et al, 2001, 2002; Gast et al, 2003; Hunter et al, 1998; Sinisterra et al, 2005). Whiteflies (Hemiptera: Aleyrodidae) aggregate in large numbers on plants (McKenzie 2002; Saeed and Samad 2017) and as thousands of excreta droplets fall from the plants during insect feeding the virus becomes aerosolized. Once the virus or bacteria becomes airborne they may travel alone or become associated with dust or other particulates in the air, and may then travel an unknown distance based upon wind flow and speed (Kormunth et al, 2018; Xie, et al, 2007). Depending upon environmental conditions such as temperature and moisture, the type of virus, or the time exposed to ultra-violet light, viruses may become inactivated within minutes to hours, or may remain viable for weeks (Marutani-Hert et al, 2009; Kanakala and Ghanim 2016). Stability traits of viruses make them detectible by molecular methods, which analyze the presence of viral nucleic acids. To use these methods the virus must first be collected. Most plant virus monitoring programs depend on taking plant tissue samples, or insects, that then undergo nucleic acid extraction and analyses with ELISA or PCR methods. Samples are usually selected based upon visual inspection by a person that will scout the field for plant tissues showing disease symptoms, or hosting insects. The use of bioaerosol sampling is based on the airflow moving the virus or bacteria through the collecting zone (Agranovski et al, 2005; Brodie et al, 2007; Mehta et al, 2000; Parvaneh, et al, 2000; Richardson et al, 2003; Yamamoto et al, 2010) (Fig. 3), thus being captured by the electrostatic field produced by the ESD.

**Figure 3.**
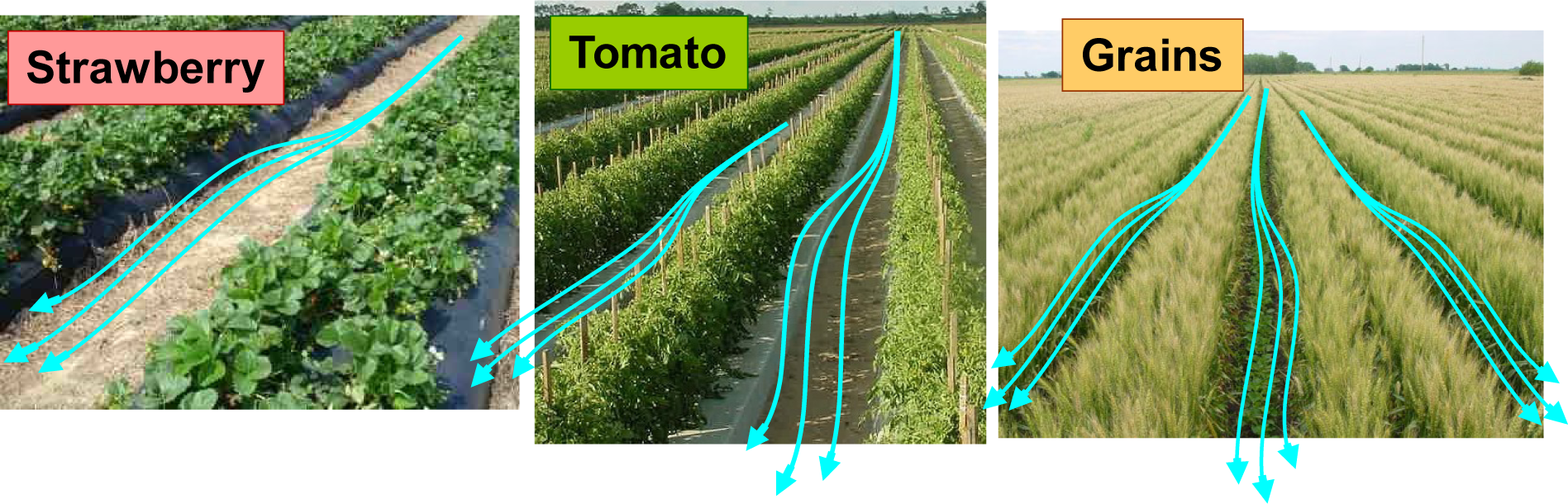
Diagrammatic representation of natural air flow through different rows of crops. As air travels down the rows it carries airborne pathogens, fungal spores and dust.

## MATERIALS AND METHODS

### Test Procedure for Sampling Poultry House Exhaust Air

The ESD was compared to settling plates and a SAS-90 (Bioscience International, Rockville, MD)(Gast et al, 2003, 2004). Previously reported results from rigorous trials with the ESD in rooms with caged chicken egg layers infected with *Salmonella enteritidis*, using selective MacConkey media (for Enterobacteriacea – gram negative bacteria), air samples were taken only at intervals of 5.6 min (500 L) and 11.1 min (1000 L) demonstrated positive collection and culturing of the bacterial pathogen (Gast et al, 2003, 2004; Richarson et al, 2003).

### Trials, Source and Maintenance of Insects and Plants

Colonies of whiteflies biotype-B [*Bemisia argentifolii*(Gennadius)] were obtained from laboratory colonies maintained by the U.S. Horticultural Research Laboratory, Ft. Pierce, FL. Whiteflies used in these experiments were maintained on dwarf cherry tomato (*Lycopersicon esculentum*cv. Florida Lanai) since 1996 by serial transfers (McKenzie 2002). Whitefly biotyping was based on RAPD PCR analysis using primers developed by De Barro and Driver (1997). Viruliferous whitefly colonies were housed in screened Plexiglass^®^ cages located in a walk-in environmental growth chamber (Environmental Growth Chambers, Chagrin Falls, OH) at 25 ± 1°C under a 16:8 light/dark photoperiod and an average light intensity of 700 μ-E PAR at top of plant canopy.

### ESD Sampling

The ESD is a multi-pin model, which runs on two nine volt batteries. The collector plate was a electically grounded metal scoopula, 17.5 cm long, 1.4 cm wide, and set ~ 2.5 cm below pins, upon which the particulates would adhere through static charge (Mitchell et al, 2003; Mitchell and Gast 2006). The ESD was set on the same table with the collector parallel to the infested plant. The ~250 adult whiteflies were given settling times of 5-6 d pre-sampling. The ESD was set up at two distances from the infested plant, at 30 cm and 60 cm, samples were collected at two time durations, 1 and 2 hrs (gels similar Fig. 2C).

### Validation of virus from whiteflies and plants

The Tomato yellow leaf curl virus (TYLCV) whitefly colony was established then serial transferred to dwarf cherry tomato cultivar. DNA was extracted from one, two, or three whiteflies using the AquaPure™ Genomic DNA isolation kit (Bio-Rad, California). DNA extraction protocol in Edwards et. al. (1991). Validation of viruses was accomplished by PCR amplification of plant or insect samples with virus specific primers). Primers: TYLCV Pico-L 5’-CGCCCGTCTCGAAGGTTC-3’, TYLCV Pico-R 5’-GCCATA TACAATAACAAGGC-3’ (Pico et al. 1998), ToMoV1: CP1 ToMoV (aka EH 289) 5’-GCC TTCTCAAACTTGCTCATTCAA T-3’, CP2 ToMoV (aka EH 290) 5’-GTTCGCAACAAA CAGAGTGTAT-3’. (Sinisterra et al, 2005; Pico et al, 1998). The PCR reaction was accomplished as follows: 45 µL Platinum^®^ PCR Supermix (Invitrogen), 2 µL Depc H_2_O, 2 µL Primers, 1 µL DNA in a final volume of 50 µL. Cycle reaction was 94° C for 2 min, then 35 cycles of (94°C for 15 sec, 46.5°C for 1:35 min, 72°C for 1:00 min).

### Begomovirus Immuno-labeling

Live adult whiteflies were immobilized by placing them in a standard −20° C freezer, for ~3-5 min, then placing the live, immobilized insect onto a drop of clear fingernail polish. Whiteflies were covered with 10 mM phosphate buffer saline (PBS), pH 7.4, (Sigma Co., P-3813, St. Louis, MO) and dissected. More details on dissection, labeling and processing described in (Hunter et al, 1998; Sinisterra et al, 2005; Cancino et al, 1995) (Fig. 1).

### Experimental design

The ESD was set at three distances (next to plant, 30 cm, and 60cm) from whitefly infested, caged tomato plants, which had been validated as being infected with TYLCV and which were contained within a growth room. The ESD was let run for two time periods, 1 hr and 2 hrs, to evaluate efficiency of virus capture. All experiments were replicated three times.

## RESULTS AND DISCUSSION

Sixteen of the 18 independent ESD samplings (88.9%) resulted in positive detection of TYLCV as a pathogen bio-aerosol being excreted by whiteflies. Twelve independent one hour sample durations resulted in 11 TYLCV positive PCR detections by the ESD collecting at both 30 and 60 cm from test plants. Six independent two hour sample durations resulted in five positive TYLCV detections at both distances. In all trials virus capture as evident by PCR band strength appeared to be greatest when closer to infested plants (Fig. 2C, other gels not shown). Our data demonstrates that Begomovirus becomes aerosolized from whitefly infested tomato, and was successfully captured by the ESD (Fig. 2C). Furthermore, the ESD is suitable to capture other aerosolized insect-transmitted viruses (Agranovski et al, 2005; Mainelis et al, 1999; Mitchell and King 1994; Mitchell and Waltman 2003) or bacteria, (Gerone et al, 1996; Holt et al, 1999; Mitchell et al, 2002), aerosolized during feeding (Suresh and Ricke 2002) (Fig. 3). However, detection does not demonstrate virus viability nor spread as a contagion. Depending upon environmental conditions such as temperature and moisture, the type of virus or bacteria, and the time exposed to ultra-violet light, microorganisms may become inactivated within minutes to hours, or remain viable for days (31, 32). The results show and we propose that electrostatic detection methods are a rapid aid in the collection of bioaerosol pathogens for detection and identification in agricultural systems (Brodie et al, 2007; Gustin et al, 2013; Kormuth, et al, 2018; Suresh and Ricke 2002; Richardson et al, 2003). The ESD will find many applications in the monitoring of different systems whether these are crop plants, or chicken houses; farm yards, or back yards, the ESD provides a low cost solution to one of the growing concerns of today’s world.

## Acknowledgements

We thank Laura Hunnicutt, Biological Science Technician, USDA, ARS, Fort Pierce, FL, for technical support and analyses.

## Disclaimer

Mention of proprietary or brand names are necessary to report factually on available data; however, the USDA neither guarantees nor warrants the standard of the product, and the use of the name by USDA implies no approval to the exclusion of others that also may be suitable. This article is in the public domain and not copyrightable. Publication may be freely reprinted with customary crediting of source.

